# Detection of neuronal OFF periods as low amplitude neural activity segments

**DOI:** 10.1101/2022.09.16.508135

**Authors:** Christian D. Harding, Mathilde C.C. Guillaumin, Lukas B. Krone, Martin C. Kahn, Cristina Blanco-Duque, Christian Mikutta, Vladyslav V. Vyazovskiy

**Affiliations:** Department of Physiology Anatomy and Genetics, Sir Jules Thorn Sleep and Circadian Neuroscience Institute, University of Oxford, Oxford, UK; The Kavli Institute for Nanoscience Discovery, Oxford, UK; Nuffield Department of Clinical Neurosciences, Sir Jules Thorn Sleep and Circadian Neuroscience Institute, University of Oxford, Oxford, UK; Institute for Neuroscience, Department of Health Sciences and Technology, ETH Zürich, Schwerzenbach, Switzerland; University Hospital of Psychiatry and Psychotherapy, University of Bern, Bern, Switzerland; Centre for Experimental Neurology, University of Bern, Bern, Switzerland; Picower Institute for Learning and Memory, Massachusetts Institute of Technology, Cambridge, USA; Department of Brain and Cognitive Sciences, Massachusetts Institute of Technology, Cambridge, USA; Translational Research Center, University Hospital of Psychiatry and Psychotherapy, University of Bern, Bern, Switzerland; Private Clinic Meiringen, Meiringen, Switzerland

## Abstract

During non-rapid eye movement sleep (NREM), synchronised neuronal activity is reflected in a specific neural oscillation observed in neocortical electrophysiological signals: a low frequency component characterised by depth-positive/surface-negative potentials known as slow waves, corresponding to alternating periods of high (ON period) and low (OFF period) spiking activity. Often overlooked in favour of slow waves, there is an interest in understanding how neuronal silencing during OFF periods leads to the generation of slow waves and whether this relationship changes between cortical layers. The foremost issue in detecting population OFF periods is the absence of a formal, widely adopted definition. Here, we grouped segments of high frequency neural activity containing spikes, recorded from the neocortex, on the basis of amplitude and asked whether the population of low amplitude (LA) segments displayed the expected characteristics of OFF periods. We corroborate previous studies showing that LA segments in neural activity signals are a uniquely identifiable structure with distinct characteristics from the surrounding signal that identify them as OFF periods including NREM sleep predominance and association with a local field potential (LFP) slow wave. In addition, we attribute new characteristics to these segments not previously associated with OFF periods: vigilance-state-dependent duration and duration-dependent homeostatic response. This could suggest that ON/OFF periods are currently underdefined, and their appearance is less binary than previously considered, instead representing a continuum.

## Introduction

Slow waves, high amplitude oscillations in the delta frequency range (0.5-4Hz) observed in electrophysiological signals are a characteristic of non-rapid eye movement sleep (NREM). A role of slow waves has been suggested in processes that are dependent on NREM sleep time, such as immune function and restoration of cognitive function (Krueger et al., 2016). Slow wave activity (SWA, spectral power in the delta frequency range) is a reliable index of sleep homeostasis (Achermann and Borbely 2011) and local SWA homeostasis is correlated with improved motor learning task performance following sleep (Huber et al., 2004). To help clarify the proposed involvement of slow waves in restorative processes and memory function, and to understand whether they are causally linked to these functions or rather simply a measurable output of an underlying process, it is vitally important that methods are developed which improve our understanding of the neurophysiology of slow waves.

Early studies, pioneered by Steriade et al. (1991), showed that slow waves during NREM reflect alternating periods of synchronised neuronal firing and silencing referred to here as ON/OFF periods (Vyazovskiy et al. 2009). It has been hypothesised that some of the functions ascribed to NREM are in fact the result of neuronal inactivity during OFF periods (Vyazovskiy and Harris 2013). For example, prolonged wakefulness activates the unfolded protein response (UPR) in mice (Naidoo et al., 2005) and exploration during wake induces double strand DNA breaks (Bellesi et al., 2016). Conversely, inactivity may provide an opportunity for rest and repair of minor cellular damage (Vyazovskiy and Harris 2013).

If it is indeed the dynamics of the synchronised neuronal activity that we are primarily interested in understanding, ON/OFF periods in local neuronal networks may provide a more direct measure of this behaviour than the slow waves it gives rise to. Despite this, far more attention has been given to detecting slow waves than ON/OFF periods. Slow wave detection methods have been developed based on amplitude and duration thresholds (McKillop et al. 2021; Mensen et al., 2016), spectral frequency (Lajnef et al., 2015; Mukovski et al., 2017) and using a neural network approach (Bukhtiyarova et al., 2016). Whilst the number and quality of events detected as slow waves vary with method and implementation (Bukhtiyarova et al., 2019), these can at least be validated against a ‘gold standard’ manual scoring by experienced observer’s using tried and test criteria. Automated slow wave detection facilitates rapid processing of large data sets. Additionally, such methods have enabled the development of real time slow wave modulation, the therapeutic applications of which are currently being explored (Geiser et al., 2020; Fehér et al., 2021).

Similarly, a range of methods have been devised to solve the problem of detecting ON/OFF period transitions. The simplest method is to apply amplitude and duration thresholds to spike trains: binary traces of high amplitude deflections (spikes) in single or multiunit activity signals (Vyazovskiy et al., 2009; Mckillop et al., 2018; Krone et al., 2021; Kahn et al., 2021; Karalis and Sirota 2022). Though widely used both for offline and online detection of ON/OFF periods, this method is sensitive to the threshold used to define spikes which is often set using visual inspection and binarization results in the loss of a large amount of data. More sophisticated methods for detecting ON/OFF periods in continuous neuronal activity signals can broadly be classified as ‘threshold-crossing’ or ‘predictive’ algorithms. ‘Threshold-crossing’ algorithms work by processing the data until a bimodal distribution is obtained upon which a threshold is applied to separate data between ON and OFF periods (Mukovski et al., 2007, De Bonis et al., 2019, Dasilva et al., 2021). ‘Predictive’ algorithms assume bimodality and assign data to one of either state based on the probability of a predictive model fitted to the data (Seamari et al., 2007; MacFarland et al., 2011; Jercog et al., 2017).

A common trend observed in the design of these detection algorithms is that they are built and tested upon a subset of data displaying clear slow waves (Mukovski et al., 2007) or ON/OFF oscillations (MacFarland et al., 2011) or acquired from anaesthetised animals where slow wave activity is generally more regular than during sleep (Mukovski et al., 2007; MacFarland et al., 2011, De Bonis et al., 2019, Seamari et al., 2007, Dasilva et al., 2021). Whilst it is likely that these methods will generalise to the case of sleep recordings, there may be advantages to designing a detection method on ‘noisy’ data characteristic of *in vivo* free-moving sleep recordings and that can be applied to the entire duration of chronic recordings used to study sleep behaviour. Another trend is to judge the performance of a detection method by comparing the output when applied to different signals recording the same neural activity (e.g. intra- vs extra-cellular, Mukovski et a., 2007) or with the output of other methods (Seamari et al., 2007; MacFarland et al., 2011). As no method can yet be called the ‘gold standard’, this makes it challenging to judge the relative merit of methods and to assess the effect of optimisation steps within the pipeline (e.g. to remove short duration state transitions/interruptions).

An alternative design approach is to detect OFF periods that match the established characteristics of slow waves during spontaneous sleep. OFF periods should occur predominantly, but not exclusively, in NREM. SWA is highest and neuronal firing rate lowest in NREM compared to wake and REM, which suggests that OFF periods are more likely to occur in this state (Vyazovskiy et al., 2009). However, slow waves are known to occur regularly during REM (Funk et al., 2016) and occasionally during both inactive and active wake (Vyazovskiy et al., 2011; Nir et al., 2011; Fisher et al., 2016). By definition, OFF periods should be consistently associated with depth positive/surface negative deflections in the electroencephalogram (EEG) corresponding to slow waves (Vyazovskiy et al., 2009), and their duration should be positively correlated with slow wave amplitude (Vyazovskiy et al., 2009). Synchronization of OFF periods between brain regions should be variable with widespread (global) OFF periods reflecting high recruitment of neurons and localised OFF periods reflecting low recruitment of neurons (Vyazovskiy et al., 2007). Finally, OFF periods should respond to changes in sleep pressure and circadian drive as a result of their homeostatic regulation (Achermann and Borbely 2011).

The aim of this study was to determine if low activity is sufficient as a criterion to detect OFF periods in neuronal signals from freely behaving mice. To achieve this we designed a simple ‘threshold-crossing’ algorithm to identify a population of low amplitude (LA) segments in high frequency neural activity informed by the distribution of spiking amplitudes during NREM sleep and assessed whether these segments recapitulated the characteristics of OFF periods expected from the latter’s association with slow waves.

## Materials and methods

All experiments were carried out in accordance with the UK Animals(Scientific Procedures) Act of 1986.

### Surgery and electrode implantation

Adult male wild type C57BL/6 mice (n=7, internally sourced from Biomedical Services at the University of Oxford, 125 ± 8 d old at baseline recording) underwent cranial surgery to record the EEG, electromyogram (EMG) and local field potential (LFP) as previously described (McKillop et al., 2018). Briefly, under ~2-3 % isoflurane anaesthesia and aseptic conditions, stainless steel screws were implanted epidurally over frontal and occipital cortical areas and referenced to a third screw implanted over the cerebellum. Stainless steel wires were implanted into the nuchal muscle to record EMG. A 16-channel laminar probe (NeuroNexus Technologies Inc., Ann Arbor, MI, USA; model: A1×16-3mm-100-703-Z16) was implanted in primary motor cortex (AP +1.1 mm; ML −1.75 mm; rotated 15° in the AP axis towards the side of the implant) to perform intracortical recordings (LFP and MUA as described in Krone et al. 2021). Each animal was also implanted with a bipolar concentric electrode (PlasticsOne Inc., Roanoke, VA, USA) in the right primary motor cortex, anterior to the frontal EEG screw in relation to a separate study (as described in Kahn et al. 2022). All electrodes were attached to an 8-pin surface mount connector (8415-SM, Pinnacle Technology Inc, KS) affixed to the skull with dental cement (Associated Dental Products Ltd, Swindon, UK).

### Electrophysiological signal acquisition

A 128-channel Neurophysiological Recording System (Tucker-Davis Technologies Inc., Alachua, FL, USA) was used to acquire tethered electrophysiological recordings. EEG and EMG signals were continuously sampled at 305 Hz and bandpass filtered between 0.1 – 100 Hz. Signals were then downsampled offline to 256 Hz via spline interpolation. Laminar probe channel signals were sampled at 25 kHz. Two signals were extracted from the laminar probe channels: decimated multiunit activity (MUA) and local field potential (LFP). Multiunit activity is the high frequency component of neural activity that contains the spiking of multiple neurons within the vicinity of an electrode. Decimation is a process for downsampling the MUA whilst retaining spiking activity by storing only the highest amplitude value, either negative or positive, recorded during a set time period. This means that if multiple neurons spike during that period, only the largest is stored, thus the majority of spikes in the decimated signal will originate from nearby neurons. The resulting signal will therefore have a high amplitude when nearby neurons are spiking and a low amplitude during periods of quiescence or when distant neurons are spiking. MUA was generated by bandpass filtering the laminar signals between 300 Hz - 5 kHz then decimating to 498Hz by splitting the signal into segments of ~50 samples and storing the maximum/minimum amplitude of alternating segments as integers. LFP was generated by zero-phase distortion bandpass filtering the laminar signal between 0.1-100 Hz and downsampling to 256 Hz via spline interpolation. All offline manipulations and analyses were performed using MATLAB (version R2020a; The MathWorks Inc, Natick, MA, USA). Prior to vigilance state scoring, signals were transformed into European Data Format as previously reported (see Fisher et al., 2016).

### Experimental design and recording procedure

For sleep recordings, animals were individually housed in sound-attenuated and light-controlled Faraday chamber cages (Campden Instruments, Loughborough, UK) with *ad libitum* food and water. A 12:12 hr light/dark cycle (lights on at 9 am = ZT0, light levels 120–180 lux) was implemented, temperature maintained at around 22 ± 2°C, and humidity kept around 50 ± 20%. Animal were given at least three days to acclimatize before two recording days starting at ZT0: a baseline day with spontaneous sleep permitted and a sleep deprivation day. On the sleep deprivation day, animals were prevented from sleeping from ZT0-ZT6 through gentle handling and the presentation of novel objects to encourage naturalistic exploration behaviour (Vyazovskiy et al., 2002).

### Vigilance state scoring and channel selection

EEG, LFP and EMG signals were used to score vigilance states in the Sleep Sign for Animals scoring environment (version 3.3.6.1602, SleepSign Kissei Comtec Co., Ltd., Nagano, Japan). Four second epochs were scored as WAKE, NREM or REM. Epochs with high frequency EEG and high amplitude EMG activity were scored as WAKE, epochs with a low frequency EEG characterised by delta band (0.5-4Hz) slow waves and sigma band (11-15Hz) spindles and a quiet EMG were scored as NREM sleep. Epochs with a wake-like EEG dominated by theta band activity and a quiet EMG were scored as REM sleep. Epochs with recording artefacts related to movement or electrostatic noise were rejected from further analyses in all channels (5.06 ± 0.19% of total recording time). Only vigilance states lasting ≥ 3 epochs were retained for further analysis to ensure clear differentiation of states. Channels with low MUA amplitude variation (i.e. without spiking activity) were rejected by visual inspection (5/112 channels). For the purpose of inter-animal comparisons, each animal was represented by a single layer 5 channel based on histology assessment of laminar probe depth (for histological methodology see Krone et al. 2021).

### Low amplitude segment extraction

The concept of distinct ON/OFF states necessitates that MUA recorded during NREM should be bimodally distributed. Assuming that OFF periods represent protracted periods of synchronised low amplitude activity, we smoothed the standardised MUA extracted from NREM sleep by convolution with a 62ms Gaussian window (width factor = 2.5, sum of weights= 1) and plotted a 1D histogram of amplitudes. This generated the bimodal distribution upon which previous ‘threshold-crossing’ algorithms have been based (Mukovski et al., 2007; De Bonis et al., 2019; Dasilva et al., 2021) (Figure 1B). However, upon comparing different channels and recording periods we found it was not always clear where the distributions diverged. As MUA amplitude during ON periods is more varied as a result of spiking events, we theorised that ON periods should be more sensitive to smoothing window length and that this property could be leveraged to facilitate differentiation of the distributions. We compared MUA amplitude smoothed with a 62ms Gaussian window (width factor = 2.5, sum of weights= 1) and a shorter 22ms Gaussian window (width factor = 2.5, sum of weights= 1) using a 2D histogram. Indeed, we consistently observed a dense region of low amplitude MUA points which we theorised may be reflecting OFF periods (Figure 1C).

**Figure 1:**
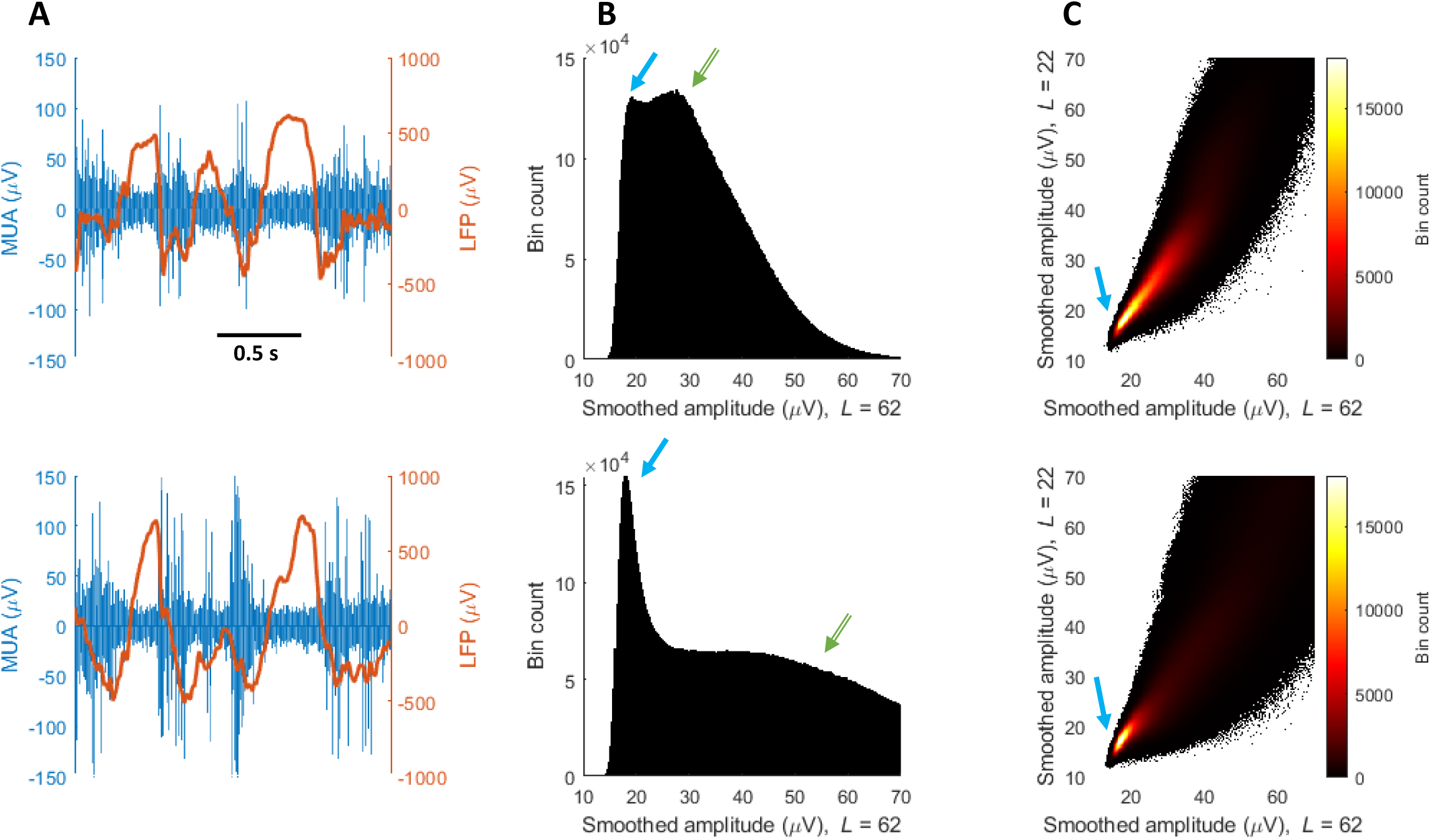
OFF period detection rationale. Top row depicts data from cortical layer 4, bottom row from layer 6. A) MUA and LFP signal from different cortical layers from the same NREM sleep interval. OFF periods can be distinguished by a reduced MUA amplitude and often by the presence of LFP slow waves. The appearance of OFF periods, in terms of amplitude and duration, differs between and within layers. B) 1D Histogram of NREM MUA amplitudes after Gaussian smoothing. L = length of smoothing window in ms, width factor = 2.5. The histogram of layer 6 has a bimodal distribution with a narrow low amplitude peak (blue arrow), which we call low amplitude (LA) data points, and a broad high amplitude peak (green arrow), which we interpret as non-LA period data points. The histogram of layer 4 is also bimodal but the peaks are closer together and have similar heights. In both cases, no obvious threshold exists at which to separate the peaks. C) 2D histogram of MUA amplitude after Gaussian smoothing with two different window lengths. The histogram is unimodal with only the low amplitude peak retained (blue arrow). Rather than setting an amplitude threshold, LA data points can now be detected by finding points belonging to this high density region.

To explore this further, we sought to find the dimensions of this low amplitude region and assign data points to using Gaussian mixture modelling (GMM). A Gaussian mixture model is a probabilistic model that assumes all the data points are generated from a mixture of a finite number of multivariate Gaussian distributions or components. Unlike k-means clustering, these components and therefore the resulting clusters do not need to be spherical in shape. To find the parameters of the Gaussian components which maximize the likelihood of the model given the data, the two-step iterative Expectation-Maximization (EM) algorithm is employed. In the expectation step, the algorithm computes posterior probabilities of component memberships for each observation given the current parameter estimates. In the maximization step, the posterior probabilities from the previous step are used to re-estimate the model parameters by applying maximum likelihood. These steps are repeated until the change in loglikelihood function is less than the tolerance (10^-5^). Once the fitted GMM has been obtained, new points can be assigned to the component yielding the highest posterior probability (hard clustering).

The primary variable that must be input for GMM is the number of components (k) to fit from the data. When using k=2 components, we unexpectedly found that the solution consistently overestimated the size of the low amplitude component. If this is indeed the data from OFF periods, this could suggest that the variation between types of ON period is greater than variation between OFF and ON periods. To resolve this, we decided to allow k to vary then select the resulting configuration that provides the optimal clustering solution. First, the smoothed MUA signals (62ms and 22ms) from a subset of NREM episodes is clustered using GMMs with k=1:8 components. Then, the optimal model is selected using a clustering evaluation index. Clustering evaluation indices are used to assess clustering performance when there is no ground truth, as is the case for binary OFF/ON period alternation. We investigated two such indices: the Calinsky-Harabasz index and the Davies-Bouldin index. The Calinsky-Harabasz index compares the dispersion within clusters with the dispersion between clusters whereas the Davies-Bouldin index compares the distance between clusters with the size of the clusters themselves. As expected, we found that the optimal clustering solution suggested by both methods produced more than 2 clusters. The lowest amplitude cluster tended to be much smaller than the equivalent with a 2-component model and more closely resembled the dense concentration of points in the MUA amplitude heatmap previously identified as the likely OFF period region (Figure 1C). In the absence of clear differences in performance, we randomly selected the Calinsky-Harabasz index as our default.

Although smoothing means that the data used for clustering is dependent on surrounding timepoints, the clustering step itself is independent of time. This contrasts with all existing descriptions of OFF periods which are understood as a feature observed in a linear time course of neuronal activity. To recapitulate the time domain, we decided to group low amplitude points into consolidated segments. First, we defined a population of time segments with below average MUA amplitude. This population was identified by taking all time segments in which the standardised MUA was below zero (Figure 2C), where the standardised MUA is calculated by taking the absolute values of the MUA then subtracting the mean of these absolute values during WAKE epochs. The waking average was chosen to represent baseline MUA so that it would not be dependent on the number and duration of OFF periods in the signal. We then isolated those which coincided with at least one time point belonging to the low amplitude cluster. This final population of segments represents the low amplitude segments used during this analysis.

**Figure 2:**
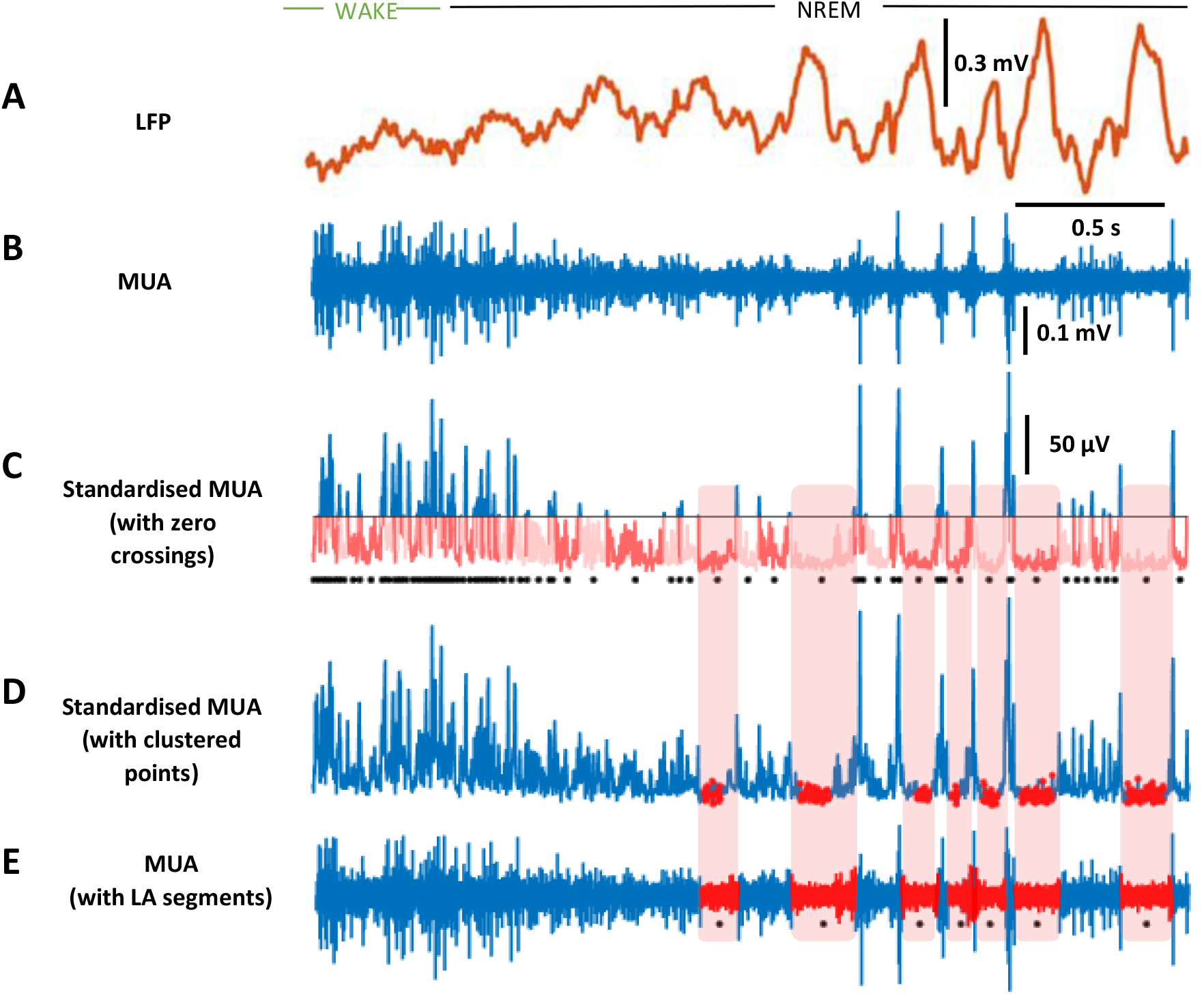
LA segment detection pipeline. Example of LA segment detection stages from a 3 second segment of layer 6 baseline day recording. A) Local field potential (LFP) trace (0.5-100Hz filtered). Note the appearance of slow waves following the transition from wake to non-REM sleep. B) Raw decimated multi unit activity (MUA). C) Standardised MUA whereby the raw signal is converted to absolute amplitude and then the mean of the absolute MUA during clean wake epochs from this 24 hour recording is subtracted. The horizontal black line marks the new mean amplitude (0μV). Each contiguous sequence of points below this line has been recoloured alternate shades of pink. These zero crossings represent the population of possible LA segments to be investigated. A black dot demarcates the centre of each zero crossing. D) Standardised MUA with low amplitude cluster points recoloured red. E) Raw MUA with final LA segments recoloured red. Final LA segments represent zero crossings which intersect with low amplitude cluster points. Black dots represent the centre of each LA segment.

Due to their independent nature, a unique clustering solution and mean MUA amplitude was generated for each channel in each animal to detect LA segments. Where the same channel was measured over multiple days, we reapplied the configuration generated from the initial day under the assumption that these signals would be dependent.

Low amplitude segment extraction pipeline:

1. Cluster MUA signal smoothed at two window lengths (62ms and 22ms, NREM only)
2. Detect lowest amplitude component
3. Assign smoothed MUA data to low amplitude cluster (all states)
4. Find negative zero-crossing half waves across all states (all states)
5. Re-assign clustered points to negative zero-crossings to find low amplitude segments

### LFP phase and channel coherence analysis

For phase analyses, LFP signal was zero-phase distortion filtered in the delta band (0.5-4Hz) to extract slow wave activity. The complex-valued analytic signal was then calculated using the Hilbert transform. The instantaneous phase angle in the interval [−*π,π*] for each element of the complex array was calculated by finding the inverse tangent and converted from radians to degrees. A phase of 0° corresponds to the peak of the oscillation and a phase of 180° to the trough of the oscillation. To measure the temporal coherence of LA segments between pairs of channels we generated the following statistic:

Pairwise channel coherence = (intercept(A|B)/sum(B) + intercept (B|A)/sum(A))/2

Where A and B are the time points of MUA signal within LA segments for two unique channels. A value of 0 denotes a situation in which all LA segments in both channels occur independently and a value of 1 denotes a situation in which all LA segments in both channels occur simultaneously. To estimate the random chance coherence between two channels we generated a randomly shuffled surrogate signal for each channel with an identical distribution of LA and non-LA segments. To achieve this, we fit a normally smoothed Kernel distribution to the distribution of LA and non-LA segment durations for each channel during the first hour of NREM sleep. We then randomly sampled these distributions in an alternating fashion to generate a shuffled sequence of LA and non-LA segments one hour in length.

### Statistics

Statistical analyses were performed in MATLAB and R. Values are reported as mean ± standard error (SEM). The normality assumption of underlying distributions was assessed for each factor level by computing a Shapiro-Wilks test. Unless stated otherwise, significance of effects was tested using one- or two-way repeated measure ANOVAs (within-subject factors “Vigilance state [NREM, REM, WAKE]”, “Day [baseline, SD]” and/or “Time”) with animal ID as a factor followed by post-hoc pairwise t-tests with Bonferroni correction. Circular statistics for phase analysis were performed using the CircStat toolbox (Berens 2009). Non-uniformity of each distribution against the von Mises distribution was confirmed using a Rayleigh test for circular data. Differences in mean direction were tested using a parametric Watson-Williams multi-sample test for equal means with Bonferroni correction. Statistical significance in all tests was considered as p<0.05. For box plots, the middle, bottom, and top lines correspond to the median, bottom, and top quartile, and whiskers to lower and upper extremes minus bottom quartile and top quartile, respectively.

### GUI design

The LA segment detection algorithm was incorporated into a MATLAB program with a user-friendly GUI that allows the detection of LA segments in new data sets, provides a visual representation of the results and generates a range of useful summary statistics (https://github.com/sjoh4302/OFFAD). Furthermore, this GUI allows for post-processing of LA segments, such as removal of brief interruptions, to fit individual user expectations of OFF periods.

## Results

### Temporal features and state dependency

We first looked at whether LA segments are more common in NREM sleep than other vigilance states. The majority of LA segments were detected in NREM sleep (91.70 ± 0.55%) and REM sleep (7.11 ± 0.55%) with a small proportion detected in WAKE (1.20 ± 0.13%) (Figure 3B). LA segment onset in all states is associated with a characteristic sharp drop in MUA amplitude (Figure 3B). There was a significant effect of state on LA segment incidence (Figure 3A, One-Way repeated measures ANOVA, F(2,12) = 68.92, p < 0.05) with LA segments occurring most frequently in NREM sleep and least frequently in WAKE. There was also a significant effect of state on the proportion of 4-second epochs containing at least one LA segment (Figure 3D, One-Way repeated measures ANOVA, F(1.02,6.11)=286.79, p < 0.05) with the vast majority of NREM epochs, the occasional WAKE epoch and over half of REM epochs (61.19 ± 1.87%) containing an LA segment. These findings show that LA segments are preferentially, but not exclusively, associated with NREM sleep as has previously been reported for OFF periods (Vyazovskiy et al., 2009, 2011; Nir et al., 2011; Funk et al., 2016; Fisher et al., 2016). We then looked at the duration of LA segments to see how this compared with previously reported OFF period descriptions. LA segment durations ranged from 8ms to greater than 1s but on average lasted 116.26 ± 2.16ms. This is slightly shorter than a previous description of OFF periods in mice of a comparable age (134ms, Mckillop et al., 2018), however this was not unexpected considering the comparison is with a detection method targeting the long duration OFF periods. Thus, this finding does not preclude the possibility that LA segments are OFF periods. The distribution of LA segments was strongly positively skewed during all states (1.54 ± 0.18, Figure 3C). The distributions were leptokurtic for WAKE (kurtosis=5.16) and REM (3.68) LA segments, such that long LA segments occurred more frequently than in a standard Gaussian distribution, but there was no excess kurtosis in the NREM distribution (3.00). Long LA segments were more common in NREM than either REM or WAKE states in which LA segment durations were similar (Figure 3E, One-Way repeated measures ANOVA, F(2,12)=34.48, p < 0.05). Whilst the relative duration of OFF periods in different states has not been reported, this does mirror the finding that slow waves in REM tend to be smaller than those occurring in NREM (Funk et al., 2016).

**Figure 3:**
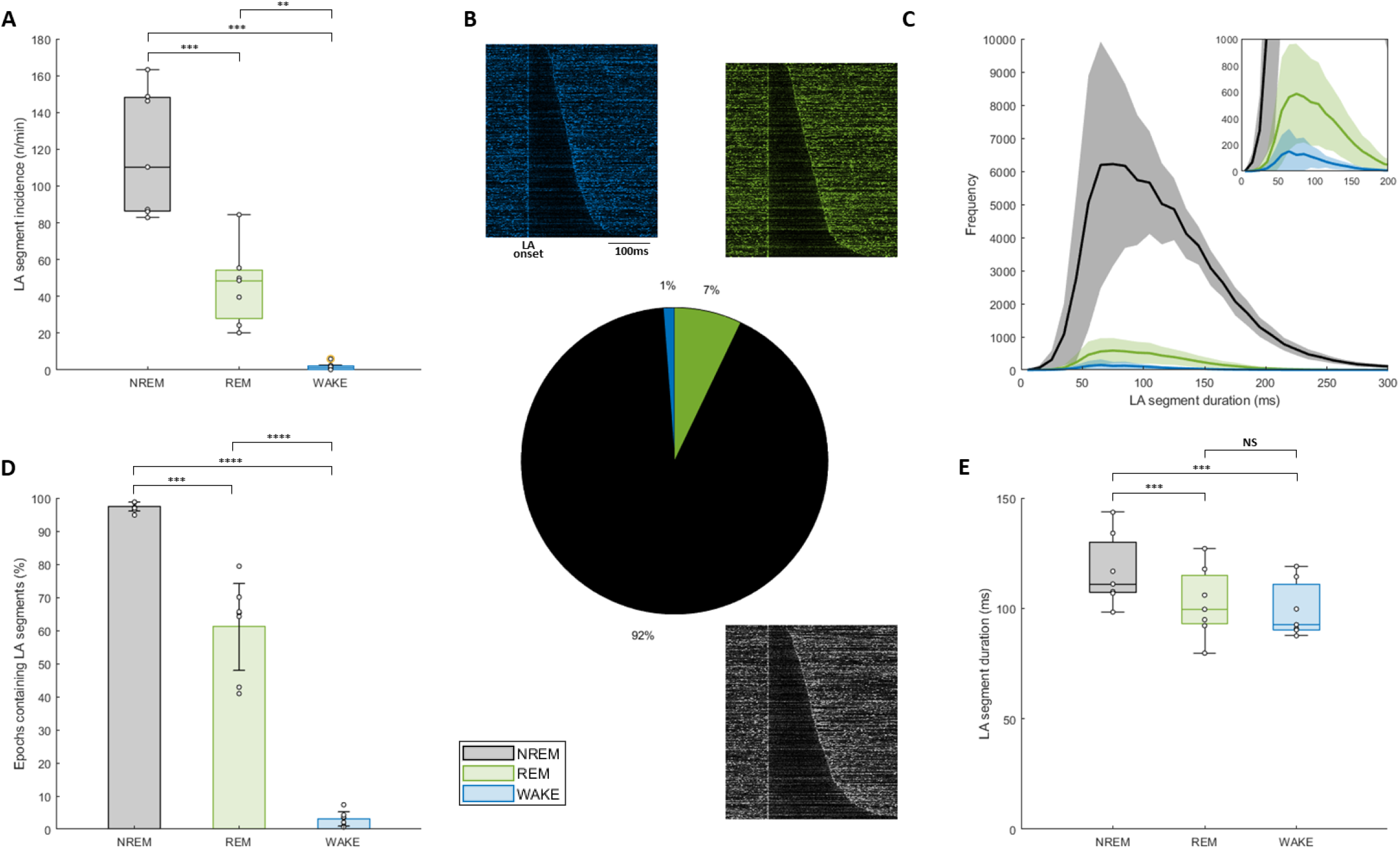
Temporal features of LA segments. A) The effect of vigilance state on the incidence of LA segments. Incidence values reported as number per minute of vigilance state. B) Global distribution of LA segments between vigilance states across study with examples of MUA amplitude during LA segments displayed for each state. Each figure shows 400 randomly selected LA segments sorted by duration. Segment time range = onset-100ms to onset+300ms. Colour of each segment scaled to MUA amplitude (dark=low, colour=high). C) Frequency of LA segments as a function of duration for each vigilance state. Mean ± SEM. Inset y-axis scaled to increase resolution of REM and WAKE states. D) The effect of vigilance state on the proportion of 4-second epochs containing an LA segment. E) The effect of vigilance state on the duration of LA segments. Black/grey=NREM, green=REM, blue=WAKE. N=7. Significance of effects assessed using one-way repeated measures ANOVA followed by post-hoc pairwise t-tests with Bonferroni correction (*P < 0.05; **P < 0.01; ***P < 0.001; ****P < 0.0001).

### Association with LFP

Evidence suggests that OFF periods coincide with LFP slow waves (Steriade et al., 1991; Vyazovskiy et al. 2009). If LA segments represent OFF periods, we would expect a similar association. We first extracted a 400ms window of LFP for each LA segment during baseline sleep from onset-100ms to onset+300ms and calculated the average LFP signal during LA segments during each vigilance state (Figure 4A). In each state, LA segments were associated with a positive deflection of the LFP which coincides with segment onset. This deflection lasts around 150ms, which if considered a half wave suggests a frequency of ~3Hz, within the delta frequency range. This deflection was bounded by slight negative deflections. These features are consistent with LA segments being time locked to LFP slow waves. To confirm this, we performed a phase analysis to determine the preferred phase of LA segment onset and exit in the delta frequency range of the LFP (Figure 4B-C). LA segment onset preferentially occurred at ~310° (NREM: 306.26 ± 1.49°, REM: 315.31 ± 1.60°, WAKE: 316.84 ± 1.52°), coinciding with the rising limb of delta oscillations. LA segment exit preferentially occurred at ~50° (NREM: 53.50 ± 0.97°, REM: 50.60 ± 1.24°, WAKE: 42.17 ± 1.43°), coinciding with the falling of delta oscillations. In all states, onset and exit phase distributions were significantly different (NREM: Watson-Williams, F(1,12)= 487.39, p<0.05, REM: Watson-Williams, F(1,12)= 286.79, p<0.05, WAKE: Watson-Williams, F(1,12)= 238.83, p<0.05).

**Figure 4:**
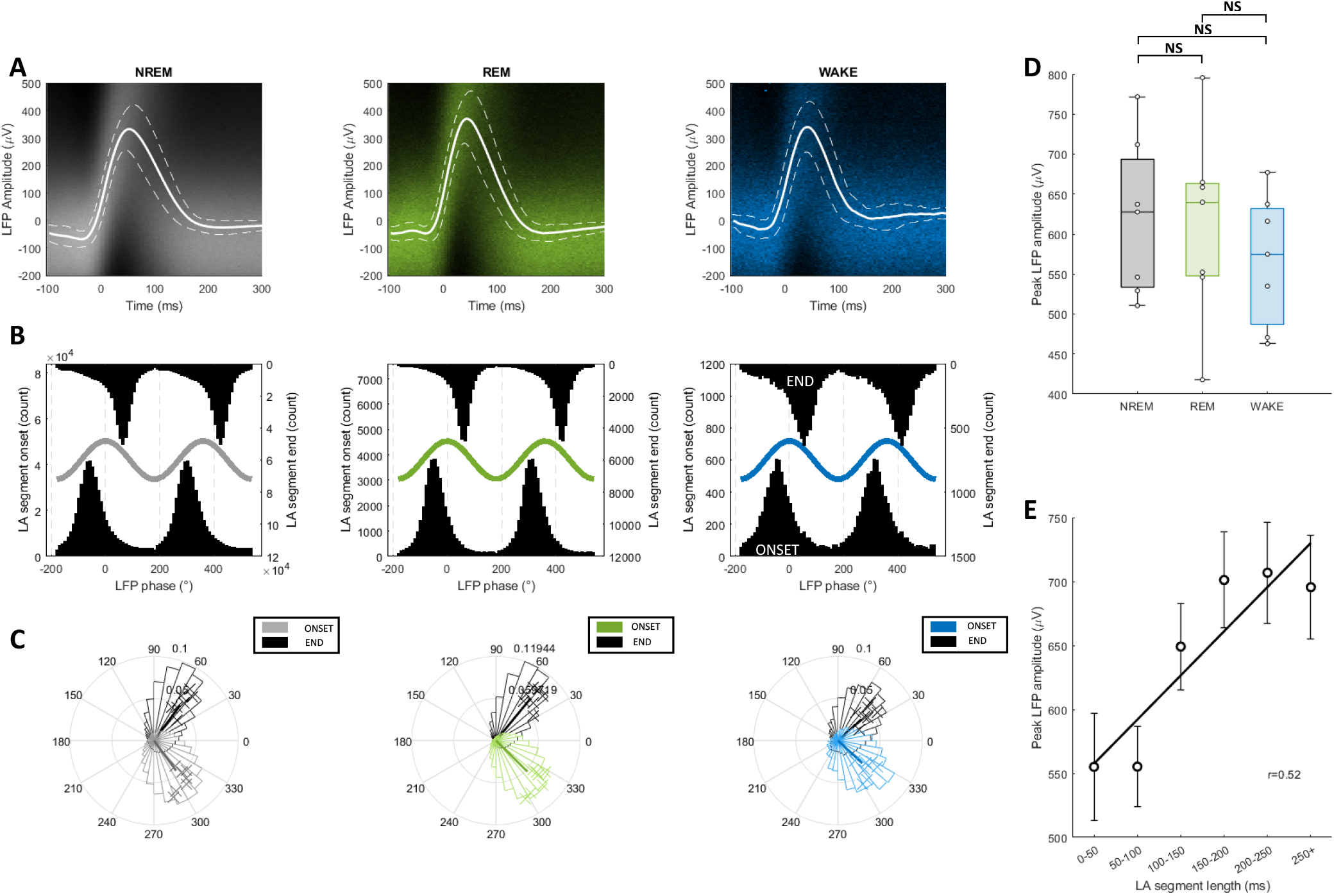
MUA LA segments associated with slow-wave activity in LFP. A) Layer 5 LFP corresponding with LA segments in each vigilance state. Composite image of LFP traces converted to heatmap (higher colour saturation = higher density) overlaid with mean ± SEM. Segment time range = onset-100ms to onset+300ms. B) Global distributions of the LFP (2-6Hz) phase corresponding to LA segment ONSET (bottom) and END (top) in each vigilance state. Example of a sinusoid wave overlaid for visualization purposes only. C) Distribution of preferred LFP (2-6Hz) phase corresponding to LA segment ONSET and END in each vigilance state (proportion) with mean resultant vector. D) Effect of state on peak amplitude in LFP corresponding to LA segments. E) Relationship between LA segment duration and peak amplitude of corresponding LFP. Least-squares regression line and significant Pearson correlation coefficient shown. Black/grey=NREM, green=REM, blue=WAKE. N=7. Mean ± SEM. Significance of effects assessed using one-way repeated measures ANOVA followed by post-hoc pairwise t-tests with Bonferroni correction (*P < 0.05; **P < 0.01; ***P < 0.001; ****P < 0.0001).

Finally, we looked at how time locked peak LFP amplitude changed with vigilance state or LA segment duration. We found a significant effect of vigilance state on LA segment duration (F(2,12)= 3.97, p = 0.047) however post-hoc pairwise t-tests did not reveal a significant difference in mean LFP amplitude between any states (Figure 4D). This suggests that LA segments in all states were associated with similar slow waves. It has been reported that slow wave amplitude increases as a function of OFF period duration (Vyazovskiy et al. 2007). In agreement with this, we found that LA segment duration was positively correlated with peak LFP amplitude (Figure 4E, r=0.52, p < 0.05).

### Homeostatic regulation

The build up and subsequent release of sleep pressure resulting from homeostatic regulation has two expected outcomes on slow waves and presumably OFF periods: slow wave activity should decrease over the course of the inactive period and slow wave activity should be higher during sleep after sleep deprivation (Achermann and Borbely 2011). To assess whether these features apply to LA segments, we analysed 6 hours of spontaneous activity during the second half of the light phase (ZT6 - ZT12) following 6 hours of baseline spontaneous activity (BL) or sleep deprivation (SD). There was a significant effect of prior sleep-wake history on occupancy time (Figure 5A, Two-Way repeated measures ANOVA, F(1,5) = 27.60, p<0.05) and duration (Figure 5B, Two-Way repeated measures ANOVA, F(1,4) = 48.14, p<0.05) of LA segments with post-hoc pairwise t-tests confirming a significant increase in the both metrics between ZT6-ZT8.5 after SD. There was a significant effect of zeitgeber time on occupancy time (Figure 5A, Two-Way repeated measures ANOVA, F(11,55) = 11.80, p<0.05) and duration (Figure 5B, Two-Way repeated measures ANOVA, F(11,44) = 7.04, p<0.05) of LA segments with both metrics decreasing as a function of time. Finally, there was a significant interaction between prior experience and zeitgeber time on occupancy time (Figure 5A, Two-Way repeated measures ANOVA, F(11,55) = 3.73, p<0.05) and duration (Figure 5B, Two-Way repeated measures ANOVA, F(1,44) = 6.85, p<0.05). To understand this further we performed post-hoc linear regressions for each condition separately. The regression of zeitgeber time against LA segment occupancy time was significant for the SD condition (linear regression model, F(1,82) = 71.756, p<0.05) but not for the BL condition (linear regression model, F(1,81) = 3.74, p>0.05). Similarly the regression of zeitgeber time against LA segment duration was significant for the SD condition (regression model, F(1,82) = 23.91, p<0.05) but not for the BL condition (linear regression model, F(1,80) = 2.54, p>0.05). To establish whether the absence of a homeostatic decrease in LA segment metrics for the BL condition was simply due to the animals having already paid the majority of their sleep debt by ZT6, we repeated regression analysis on data from the entire light period (ZT0-12) and indeed found significant regressions of zeitgeber time against LA segment occupancy (Supplementary Figure 1A, linear regression model, F(1,165) = 14.28, p<0.05) and duration (Supplementary Figure 1B, linear regression model, F(1,164) = 10.97, p<0.05).

**Figure 5:**
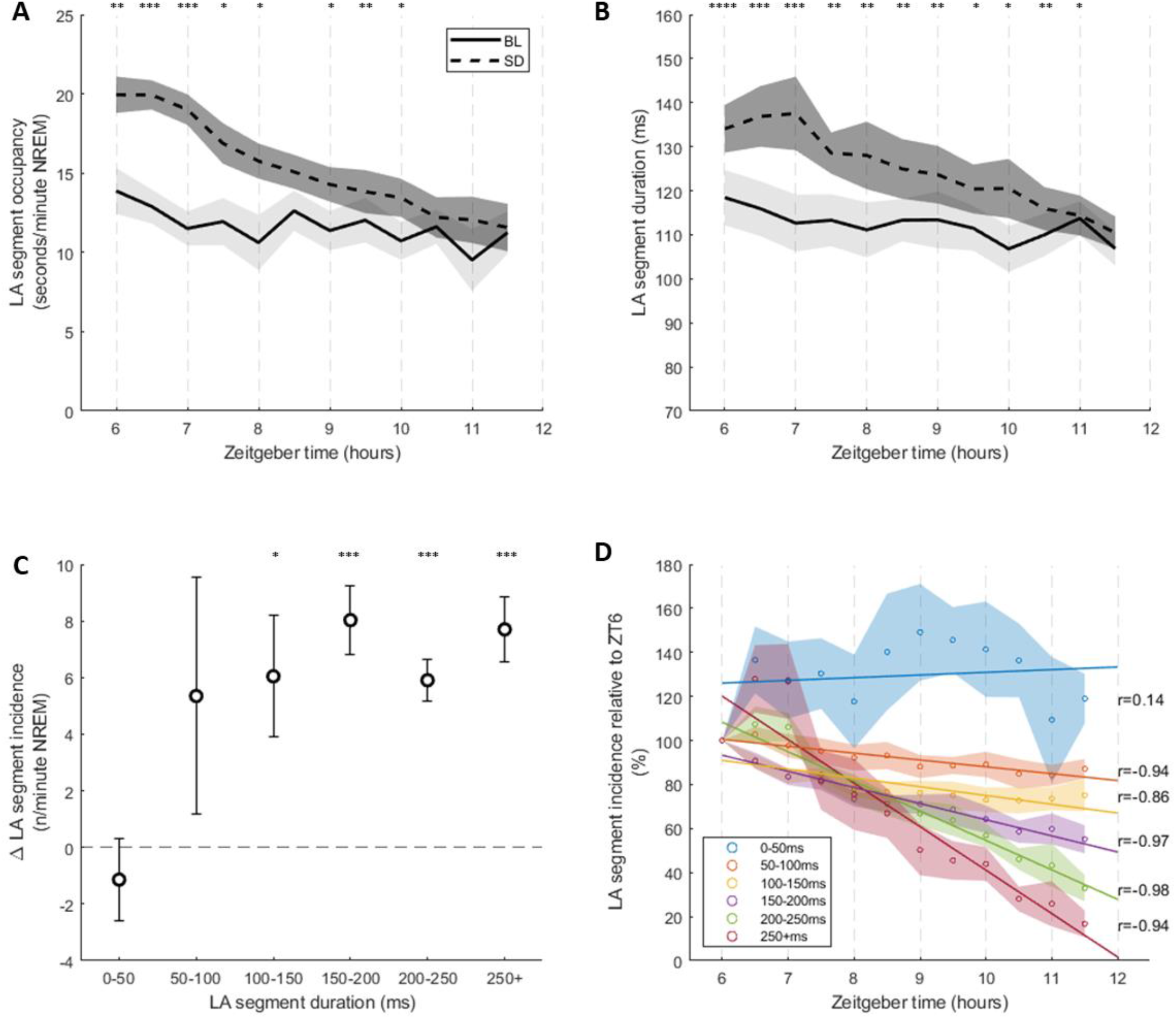
LA segments are homeostatically regulated. A) Relationship between LA segment occupancy and time during NREM sleep between ZT6 and ZT12 on baseline (BL) and sleep deprivation (SD) days (30min bins). Occupancy values reported as seconds of LA segments per minute of NREM. B) Change in LA segment duration during NREM sleep between ZT6 and ZT12 on BL and SD days (30min bins). C) Relationship between LA segment duration and the change in LA segment incidence between BL and SD days during the 1^st^ hour after sleep deprivation. A change >0 describes an increase in incidence with the sleep deprivation treatment. D) Relationship between LA segment occupancy and time during NREM sleep between ZT6 and ZT12 on the sleep deprivation day as a function of LA segment duration. Circles denote mean coherence and lines show least-square regression for each duration category. N=7^i^. Mean ± SEM. Significance of effects assessed using two-way repeated measures ANOVA followed by post-hoc pairwise t-tests with Bonferroni correction (*P < 0.05; **P < 0.01; ***P < 0.001; ****P < 0.0001). Only significant Pearson correlation coefficients shown. ^i^One animal lacked LA segments <50ms for at least one time bin so was not included in the analysis for this group of segments in D.

OFF periods are usually portrayed as an all or none phenomenon. Thus OFF period duration would not be expected to have a significant effect on a feature such as homeostatic regulation. To assess whether this is true for LA segments, we binned segments in 5 groups based on duration and looked at incidence as a marker of homeostatic regulation. We found that there was a significant increase in LA segment incidence during the first hour after sleep deprivation as compared to spontaneous activity for segments of 100-150ms (Figure 5C, Bonferroni adjusted paired T-test, t(6)= −2.81, p<0.05), 150-200ms (Figure 5C, Bonferroni adjusted paired T-test, t(6)= −6.67, p<0.05), 200-250ms (Figure 5C, Bonferroni adjusted paired T-test, t(6)= −8.06, p<0.05) and 250+ms (Figure 5C, Bonferroni adjusted paired T-test, t(6)= −6.69, p<0.05) in duration. There was no significant change in incidence for LA segments <50ms or 50-100ms in duration, though incidence of 50-100ms LA segments varied considerably between subjects with 5/7 animals showing an increase of >5 segments per minute following sleep deprivation. Furthermore, whilst the incidence of LA segments >50ms in duration was negatively correlated with time (Figure 5D) in the 6 hours following sleep deprivation, there was no correlation for segments <50ms. We noted that the magnitude of relative decrease in incidence over time appeared to increase with LA segment duration (Figure 5D). To confirm this, we fit a linear regression model to the distribution of relative LA segment incidence to zeitgeber time for each LA segment duration category. Except for segments <50ms in duration, the regression of zeitgeber time against LA segment incidence was significant across all time bins (see Supplementary Figure 2). Furthermore, the estimate for the predictor variable (i.e. zeitgeber time) became increasingly negative with increasing LA segment duration (β = −3.11 (50-100ms) < −3.99 (100-150ms) < −7.33 (150-200ms), −13.41 (200-250ms) < −19.75 (250+ms)). Overall, the homeostatic response of LA segments does appear to depend on their duration, contrary to the expectation of OFF periods being an all or none phenomenon.

### Interchannel coherence

OFF periods are both a global and a local phenomenon. To investigate whether this was also true of our LA segments, we determined the temporal coherence between channels separated by different distances along the laminar probe and compared coherence at different interchannel distances (Figure 6A, Supplementary Figures 3/4). To determine whether any coherence found was greater than chance, we generated surrogate channels by randomly shuffling LA and non-LA segments from the original channels (see Methods). To avoid vigilance state-dependent effects we only used data from NREM sleep for coherence analysis and shuffling. There was a significant effect of channel type (Figure 6B, Two-Way repeated measures ANOVA, F(1,6) = 89.78, p < 0.05), which suggests the LA segments in original channels, which have a higher mean coherence across all interchannel distances, are more synchronous than LA segments in surrogate channels. This finding highlights the generalised synchronicity of OFF periods across laminar layers. Furthermore, there were significant effects of interchannel distance (Figure 6B, Two-Way repeated measures ANOVA, F(13,78) = 11.16, p < 0.05) and interchannel distance * type interaction (Figure 6B, Two-Way repeated measures ANOVA, F(13,78) = 20.66, p < 0.05). We used Pearson’s correlation to interrogate the interaction term further and found that coherence was negatively correlated with original interchannel distance (Figure 6B, r=-0.45, p<0.05) but was not correlated with surrogate interchannel distance. This suggests that in terms of LA segment incidence, neighbouring channels are more synchronous than distant channels for original channels but not for surrogate channels where channel pairs are equally synchronous across the range of interchannel distances. This is consistent with global neocortical OFF periods displaying subtle layer dependent effects (i.e. localised dynamics).

**Figure 6:**
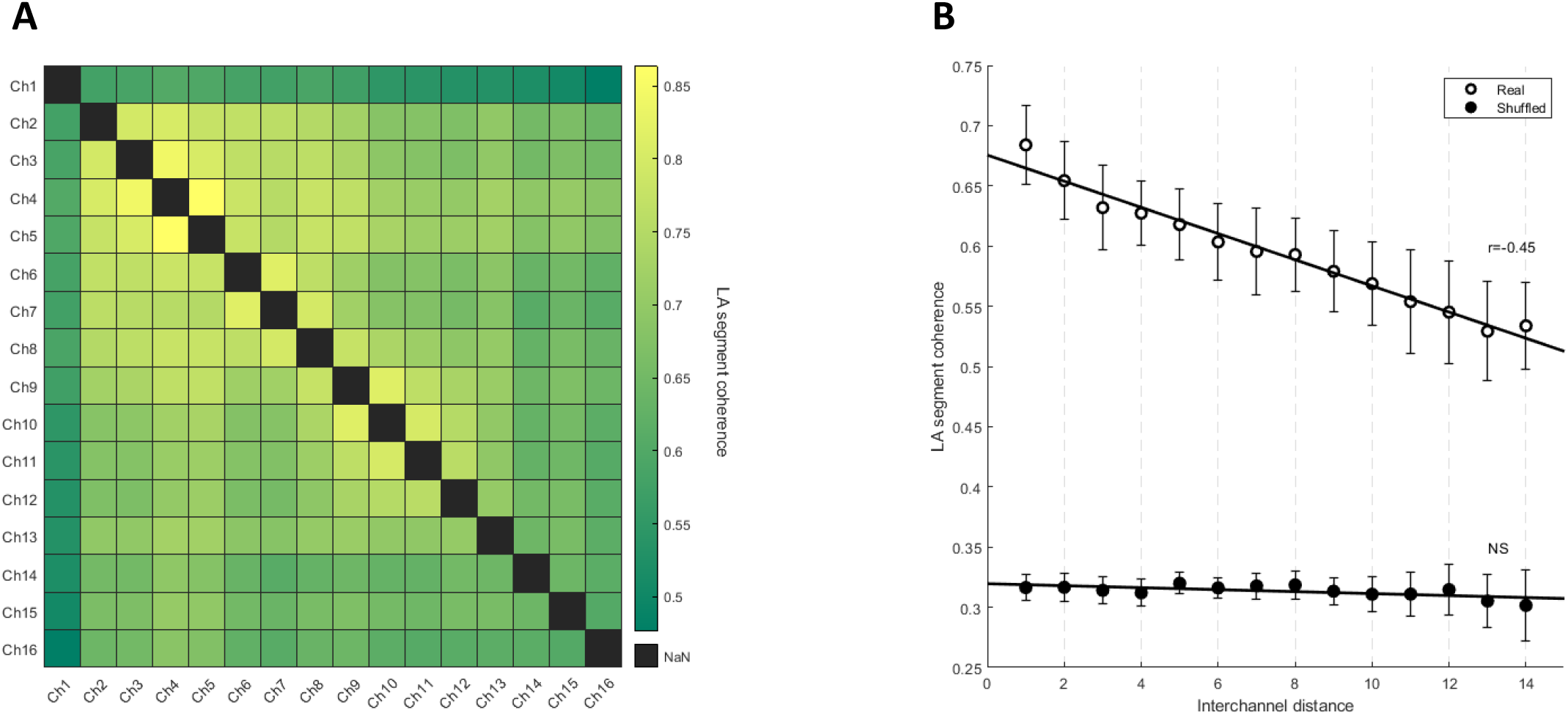
LA segments have both local and global dynamics. A) Example temporal coherence matrix between 16 channels arranged by depth for one animal (green=low coherence, yellow=high coherence). Temporal coherence reported as the average proportion of time spent in the low amplitude state during which both channels are synchronously in a low amplitude state (see Methods). B) Relationship between the temporal coherence of LA segments in pairs of channels and the distance between those channels along the laminar probe for both original and surrogate channels. N=7. Mean ± SEM. Least-squares regression lines and significant Pearson correlation coefficients shown.

## Discussion

Previous attempts to identify OFF periods in high frequency neural activity have used methods of varying complexity and levels of processing to fit their results to co-occur with slow waves. Here we show that a population of low amplitude segments can be extracted from high frequency neural activity without prior knowledge of concomitant LFP activity which fit the expected characteristics of OFF periods.

LA segments present as brief reductions in MUA amplitude, primarily during NREM but also appear in REM and WAKE. LA segments usually last approximately 100ms but duration is variable with many segments upwards of 1s occurring. LA segments are associated with a positive deflection in deep neocortical LFP (layer 5) characteristic of slow waves. Furthermore, LA segment onset and exit times are phase locked to the LFP in the delta range where slow waves occur. The duration and total occupancy time per minute of NREM sleep of LA segments decreases throughout the inactive period of mice during spontaneous activity (Supplementary Figure 1) and after sleep deprivation (Figure 5) whilst the absolute magnitude of both metrics is increased by sleep deprivation, consistent with expectations of a phenomenon regulated by sleep homeostasis (Achermann and Borbely 2011). Finally, LA segments are temporally synchronised across neocortical laminar layers at both a global and local scale. Together, these findings strongly suggest that LA segments represent OFF periods.

This finding is important for two key reasons. First, we extend the work of previous studies showing that OFF periods can be detected based on high frequency neural activity amplitude alone without reference to slow waves (e.g. De Bonis et al., 2019) by applying this methodology to recordings from freely behaving mice and by showing that it can be leveraged to detect OFF periods in all vigilance states. This permits detection of OFF periods in cases where the LFP is uninformative and opens up the possibility of studying LFP slow waves and MUA OFF periods as separate measures of the phenomenon of synchronised neuronal silencing events, each providing unique information at different scales of integration across brain regions. The MUA-based local assessment of OFF periods will be of particular importance for advancing the understanding of layer-specific dynamics in neocortex because LFP signals are influenced by volume conductance from adjacent layers and neighbouring brain regions. Second, we provide a simple method for detecting OFF periods that can be implemented as part of an easily accessible toolbox. This will allow for greater accessibility of OFF period analyses in electrophysiological sleep research and will help foster the growing interest in ‘local sleep’ effects in behavioural neuroscience.

### Homeostatic regulation of LA segments depends on segment duration

Unexpectedly, we found that the duration of LA segments had an effect on their homeostatic response. LA segments <100ms in duration showed a different homeostatic response to segments >100ms in duration. LA segments of 50-100ms occurred more frequently immediately following sleep deprivation than after a few hours of recovery sleep, consistent with homeostatic regulation. However, they showed no increase in incidence after sleep deprivation compared with the same subjective hour on a day without prior sleep deprivation. LA segments <50ms showed neither a homeostatic decrease in incidence after sleep nor an increase after sleep deprivation. Most importantly, longer segments showed a steeper decrease in incidence following sleep deprivation than shorter segments, evidence of a greater response to a build-up of sleep pressure.

One explanation of these results is that short LA segments do not represent OFF periods as they are classically described. This may best explain the absence of a clear homeostatic response in the shortest LA segments (<50ms). Therefore, we suggest that these short LA segments are not consistent with current descriptions of OFF periods and that, as others have done previously (Jercog et al., 2017), a minimum duration of 50ms should be introduced to remove brief decreases in MUA amplitude that do not behave in a similar way to longer decreases commonly identified as OFF periods. However, this explanation does not account for the mixed results for 50-100ms LA segments and the graded intra-day homeostatic response of LA segments by duration. Two hypotheses would explain this result. First, the proportion of LA segments that represent functionally relevant OFF periods as opposed to transient decreases in MUA amplitude may increase with OFF period duration. As such, the longest LA segments may show the strongest homeostatic response as the effect is less diluted by non-homeostatically regulated noise. Second, our findings are evidence that OFF periods and their associated dynamics lie on a continuum depending on the strength of neuronal recruitment and therefore duration. Greater recruitment leads to longer OFF periods which display a stronger homeostatic response to sleep pressure, either spontaneously generated or via sleep deprivation. We recognise that both hypotheses may fit the data, however we suggest that the latter may be more likely considering the abundant evidence of other OFF period-like behaviour for LA segments.

#### LA segment characteristics show state dependency

As part of our validation of LA segments as OFF periods, we chose to differentiate between LA segments occurring in different vigilance states. As expected, LA segments occurred most frequently in NREM but also occurred in the majority of REM epochs and occasionally during WAKE. Furthermore, LA segments had a similar LFP profile independent of vigilance states suggesting all were associated with slow waves. Less expected was the finding that LA segments during NREM were longer than LA segments in REM and WAKE. This could suggest that OFF periods during WAKE and REM do not achieve the same recruitment of neurons to the OFF state than in NREM. WAKE and REM OFF periods may be more localised as WAKE and REM are more ‘active’ states and as such the neuromodulatory milieu disrupts the formation of synchronous activity.

### Conclusion

We provide strong evidence that OFF periods can be detected by clustering together cortical multiunit activity segments of similarly low amplitude of extracellularly recorded neuronal spiking. These low amplitude segments show many characteristics expected of OFF periods, including NREM predominance, a strong association with LFP slow waves, sleep homeostasis and temporal coherence across cortical layers. Furthermore, we find that the incidence of longer LA segments respond more strongly to sleep pressure than short LA segments and that LA segments are longer in NREM than REM or WAKE. These vigilance-state- and duration-dependent effects were not previously described for OFF periods but these findings may represent additional OFF period features that have either been overlooked or are only revealed with multiunit activity amplitude-only detection methods.

---

**Supplementary Figure 1:**
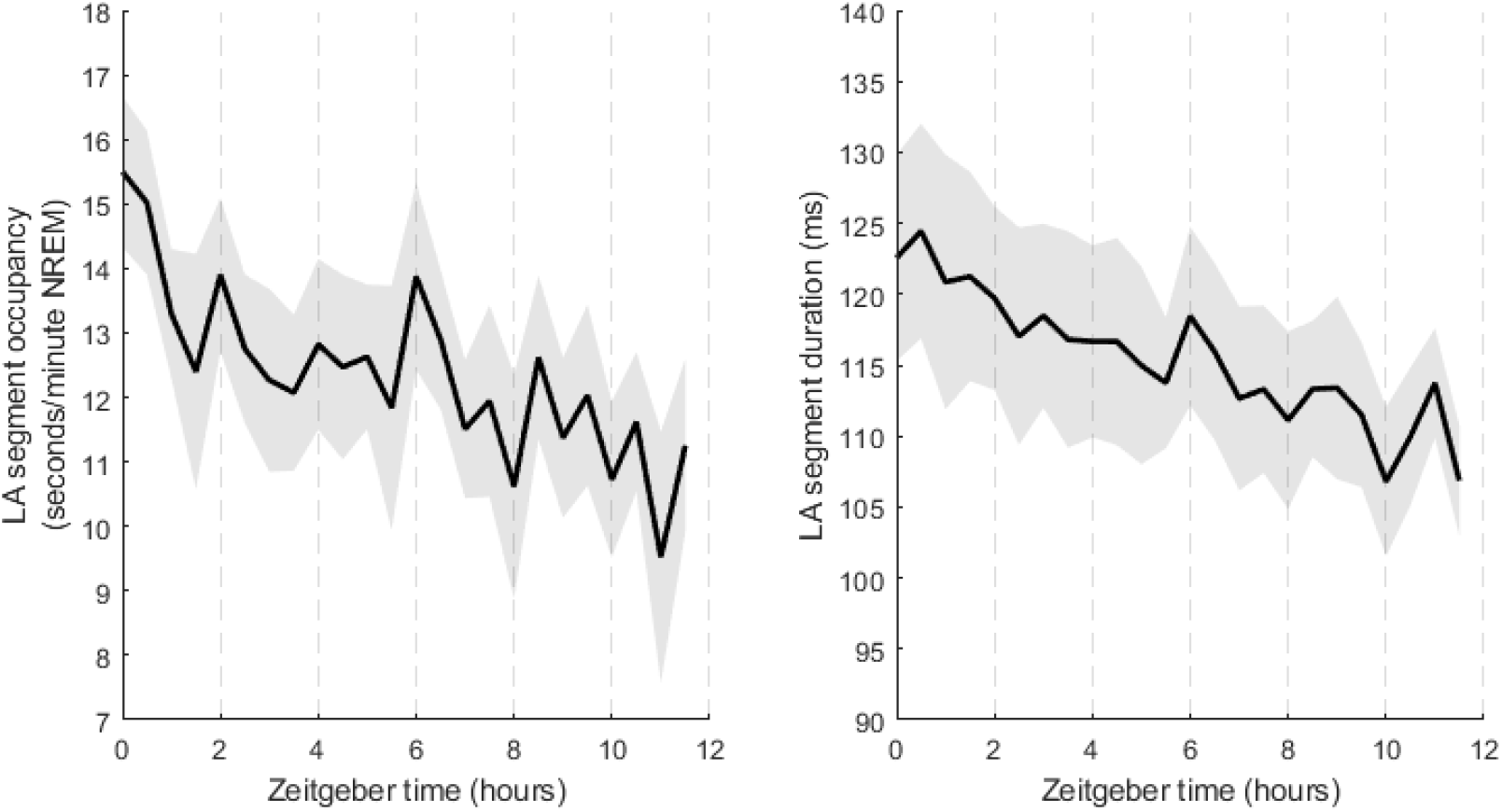
LA segment occupancy (A) and duration (B) on baseline day across light period (ZT0-ZT12). N=7. Mean ± SEM.

**Supplementary Figure 2:**
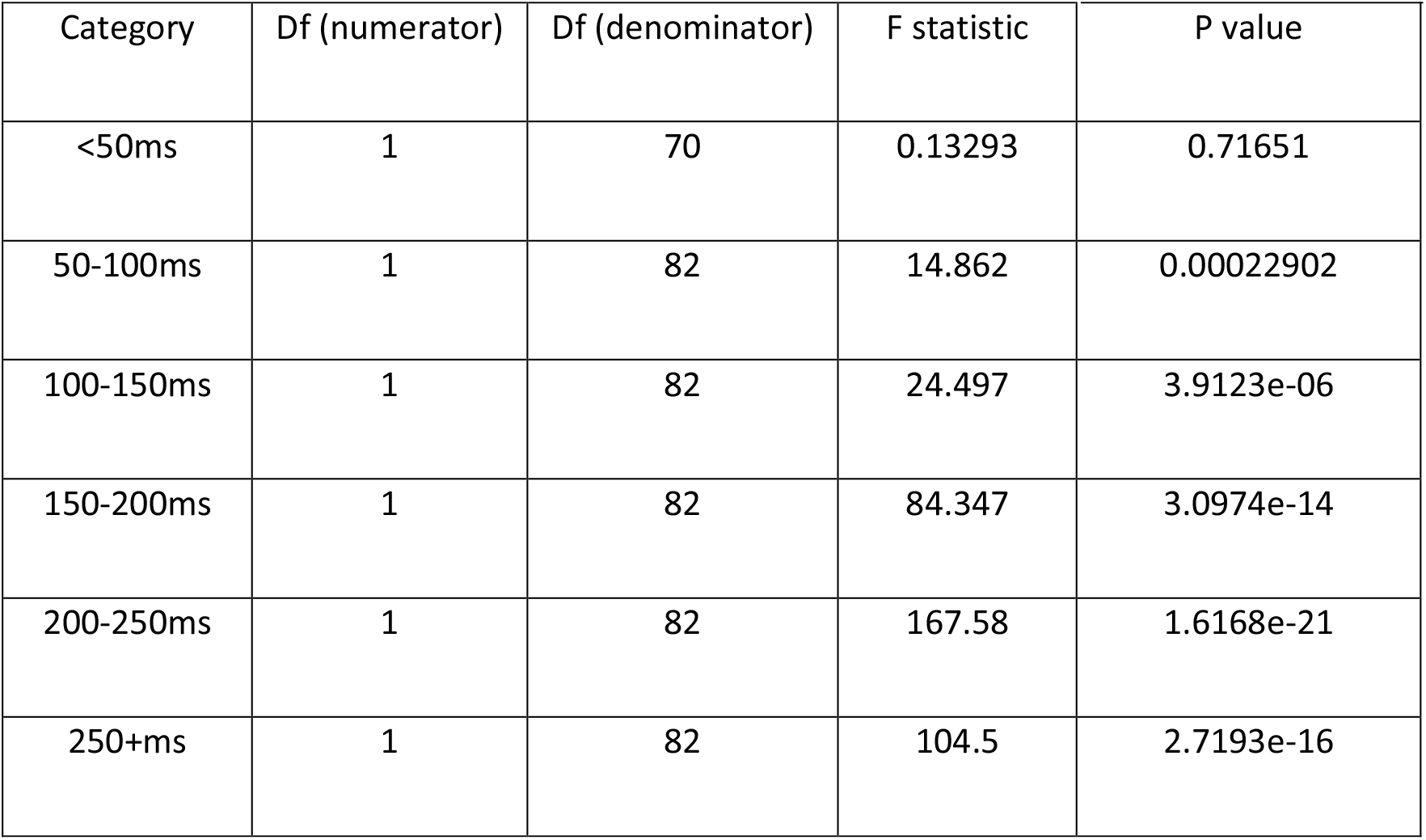
Summary statistics for linear regression of LA incidence (relative to ZT 6) against zeitgeber time for each LA segment duration category. Df = degrees of freedom.

**Supplementary Figure 3:**
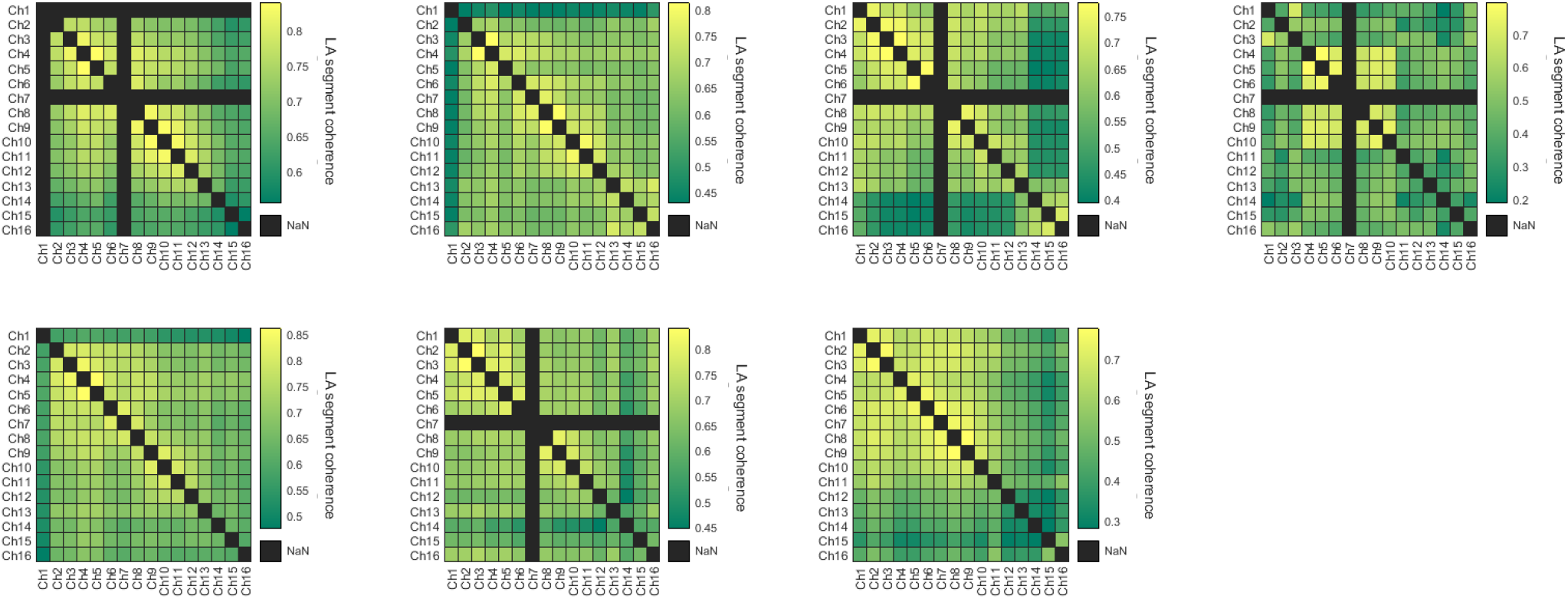
All original channel coherence matrices (N=7).

**Supplementary Figure 4:**
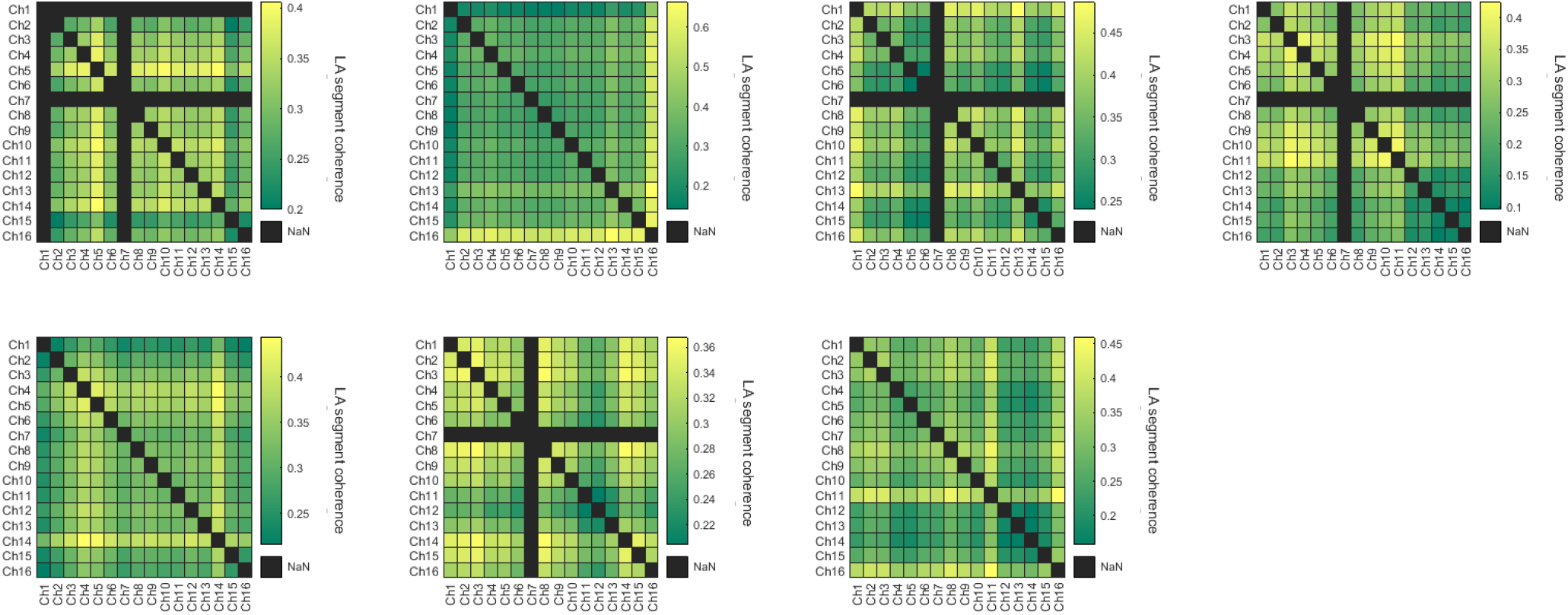
All surrogate channel coherence matrices (N=7).

## Acknowledgements

LBK was supported by a Wellcome Trust PhD studentship (203971/Z/16/Z) and by a Mann Senior Scholarship in medical sciences at Hertford College, Oxford. CDH was supported by funding from the Engineering and Physical Sciences Research Council (EPSRC, EP/S515541/1). VVV is supported by Medical Research Council (UK) grant MR/S01134X/1.

## Author contributions

CDH, MCCG, LBK and VVV designed the study. CDH analysed the data and developed the Matlab GUI. MCK, LBK, CBD and MCCG conducted the experiments. LBK and MCK performed histology. CDH and VVV wrote the manuscript with input from all authors.

## Data and code availability statement

Code for OFF period detection (OFFAD) is available on GitHub (https://github.com/sjoh4302/OFFAD).

## Competing interests

The authors declare no competing interests.

## Notes

### Competing Interest Statement

The authors have declared no competing interest.

